# SARS-CoV-2 comprehensive receptor profiling: mechanistic insight to drive new therapeutic strategies

**DOI:** 10.1101/2021.03.11.434937

**Authors:** Sarah MV Brockbank, Jo Soden, Raquel Faba-Rodriguez, Lyn Rosenbrier Ribeiro, Catherine Geh, Helen Thomas, Jenni Delight, Lucy Coverley, W Mark Abbott, Jim Freeth

## Abstract

Here we describe a hypothesis free approach to screen for interactions of SARS-CoV-2 spike (S) protein with human cell surface receptors. We used a library screening approach to detect binding interactions across one of the largest known panels of membrane-bound and soluble receptors, comprising 5845 targets, expressed recombinantly in human cells. We were able confirm and replicate SARS-CoV-2 binding to ACE2 and other putative coreceptors such as CD209 and CLEC4M. More significantly, we identified interactions with a number of novel SARS-CoV-2 S binding proteins. Three of these novel receptors, NID1, CNTN1 and APOA4 were specific to SARS-CoV-2, and not SARS-COV, with APOA4 binding the S-protein with equal affinity as ACE2. With this knowledge we may further understand the disease pathogenesis of COVID-19 patients and how infection by SARS-CoV-2 may lead to differences in pathology in specific organs or indeed the virulence observed in different ethnicities. Importantly we illustrate a methodology which can be used for rapid, unbiassed identification of cell surface receptors, to support drug screening and drug repurposing approaches for this and future pandemics.

## INTRODUCTION

Over the past two decades there have been 3 major epidemics posing risk to human life, all of which have been caused by strains of viral betacoronaviruses; the Middle East Respiratory Syndrome coronavirus (MERS-CoV), Severe Acute Respiratory Syndrome coronavirus (SARS-CoV) and the COVID-19 coronavirus pandemic (SARS-CoV-2), which has seen the loss of >2.6 million lives worldwide (as of March 2021). When a new previously unidentified virus emerges, a lack of knowledge of the virus and understanding of the mechanism by which it infects people poses a major challenge with respect to applications of current treatment and to the design and development of potential novel targeted therapies. As such identification and access to relevant and rapid technologies that can aid deconvolution of the viral life cycle to inform therapeutic options is essential.

Viral infections are initiated when virus particles bind to host cellular receptors, mediating viral entry, hijacking the host cellular machinery for replicating the viral DNA and driving the manifestation of infectious disease. Betacoronaviruses all share a similar structural configuration with a central body core, containing the viral DNA, decorated with cell surface spike glycoproteins (S), giving it the distinct crown-like appearance. The spike proteins consist of an S1 domain responsible for host receptor binding, and an S2 domain responsible for cell membrane fusion, important aspects for determining host tropism and transmission capacity (Li et al, 2016; Lu et al, 2015; Wang et al, 2016; He et al, 2004). In addition, post-translational modifications, such as glycosylation, may extend the diversity of the interaction of the spike protein with other proteins, adding to the complexity and specificity of the mechanism by which each virus causes infection (Fung et al, 2018).

Across the human betacoronoviruses, SARS-CoV-2 shares 86.22% overall homology with SARS-CoV and 45.57% with MERS-CoV (Lu et al, 2020; Jaimes et al, 2020). However, comparing commonality of the spike region alone, using the EMBOSS Needle pairwise sequence alignment tool, there is lower sequence identity with just 76.4% between SARS-CoV S and SARS-CoV-2 S, with the most variable region occurring within the S1 domain. Given the variation between the betacoronaviruses in sequence identity and post translational modification of the spike proteins, differences in receptor recognition, mechanisms of viral entry and tropism between the strains are expected.

Angiotensin converting enzyme 2 (ACE2) was identified as the host-recognition receptor for viral entry of SARS-CoV (Li et al, 2003; Wang et al, 2008). Despite the differences in amino acid similarity within the receptor binding domain, SARS-CoV-2 has also been shown to use ACE2 for viral entry (Zhao et al, 2020; Hoffmann et al, 2020) and to bind with even higher affinity than SARS-CoV (Tai et al 2020; Walls et al 2020). ACE2 has a wide distribution across human tissues, including lung, liver, stomach, ileum, colon and kidney (Zou et al, 2020), correlating with clinical effects in these target organs. However, there is a discordance in some tissues between expression levels of ACE2 and the level of SARS-CoV-2 infection, suggesting SARS-CoV-2 may depend on a co-receptor or other auxiliary membrane proteins to facilitate its infection and tropism in these tissues (Trypsteen et al, 2020).

Several groups have conducted studies to identify receptors and co-receptors involved in mechanisms of infection for SARS-CoV-2. Qi et al (2020) looked at co expression of receptors where those with similar expression patterns to ACE2 were considered to be putative coreceptors e.g. ANPEP, DPP4 and ENPEP; Wei et al (2020) screened a genome-wide pooled CRISPR library for pro and anti-viral responses to SARS-CoV-2 infection; and Gordon et al (2020) mapped protein-protein interactions between human and SARS-CoV-2 proteins using affinity purification mass spectrometry (Gordon et al, 2020). Gu et al (2020) carried out an analysis of SARS-CoV-2 spike interactions with the host receptome and identified 12 putative receptors with KREMEN1, ASGR1 and CLEC4M showing greatest affinity. A number of cell surface lectins have also been identified which bind spike protein of both SARS-CoV and SARS-CoV-2. CLEC4G, CLEC4M and CD209 (CLEC4L) have been cited as receptors of SARS-CoV (Yang et al, 2004; Marzi et al, 2004; Gramberg et al, 2005; Li et al, 2008), the latter two also cited as coreceptors for SARS-CoV-2 (Brufsky & Lotze, 2020). The presence and biodistribution of these glycan specific cell surface lectins, together with host-specific viral glycosylation could influence transmission.

In an effort to rapidly develop therapies for the COVID-19 pandemic, drug repurposing has been explored with efforts directed at mechanisms of virus replication and viral entry (Khodadadi et al, 2020), but there are limited therapeutic options for inhibiting the receptors identified to date. While there are no specific inhibitors of ACE2 binding, novel receptors or coreceptors involved in SARS-CoV-2 infection may therefore offer potential as new therapeutic targets.

Here we describe a hypothesis free approach to screen for novel interactions of SARS-CoV-2 spike protein with human cell surface receptors. We designed, expressed and purified an optimal recombinant natively folded trimeric spike protein of SARS-CoV-2 and used a high throughput library screening approach to detect binding interactions. To our knowledge, this is the largest panel of membrane bound and soluble receptors, comprising 5845 targets, expressed recombinantly in human cells. We were able to confirm and replicate SAR-CoV-2 binding to ACE2 and other putative coreceptors CD209, ASGR1 (CLEC4H1) and CLEC4M. More significantly, we identified additional novel receptors, CNTN1, NID1 and APOA4, which have not been associated with coronavirus infection to date. With this knowledge we may further understand disease pathogenesis of COVID-19 patients and how infection by SARS-CoV-2 may lead to failure of specific organs or virulence in different ethnicities. Importantly we illustrate a methodology which can be used for rapid and unbiassed identification of cell surface receptors, to support drug screening and drug repurposing approaches for this and future pandemics.

## RESULTS

### Experimental design

The construct design for this study was based on published literature (Gui et al, 2017; Pallesen et al, 2017; Kirchdoerfer et al, 2018; Amanat et al, 2020; Walls et al, 2017; Wrapp et al, 2020). Constructs included a modified S1/S2 domain boundary cleavage site to avoid sample heterogeneity, as well as a pair of proline mutations to stabilise the spike proteins in the prefusion conformation (Fig 1A). Soluble, trimeric SARS-CoV-2 and SARS-CoV spike proteins fused at the C-terminus to the Fc region of human IgG as a means of detection in the screening process, and to a 6His tag (SARS-CoV-2 S and SARS-CoV S, respectively) were produced recombinantly using the HEK-293 expression system with yields of purified protein of up to 1.7 mg/L culture. Reducing SDS-PAGE analysis of the purified proteins showed single bands between 140 and 200 kDa (Fig 1B). However, fusion of the Fc region also resulted in dimerization of the spike trimers, with molecular weights estimated to be greater than 600 kDa by size exclusion chromatography analysis (Fig 1C).

**Figure 1.**
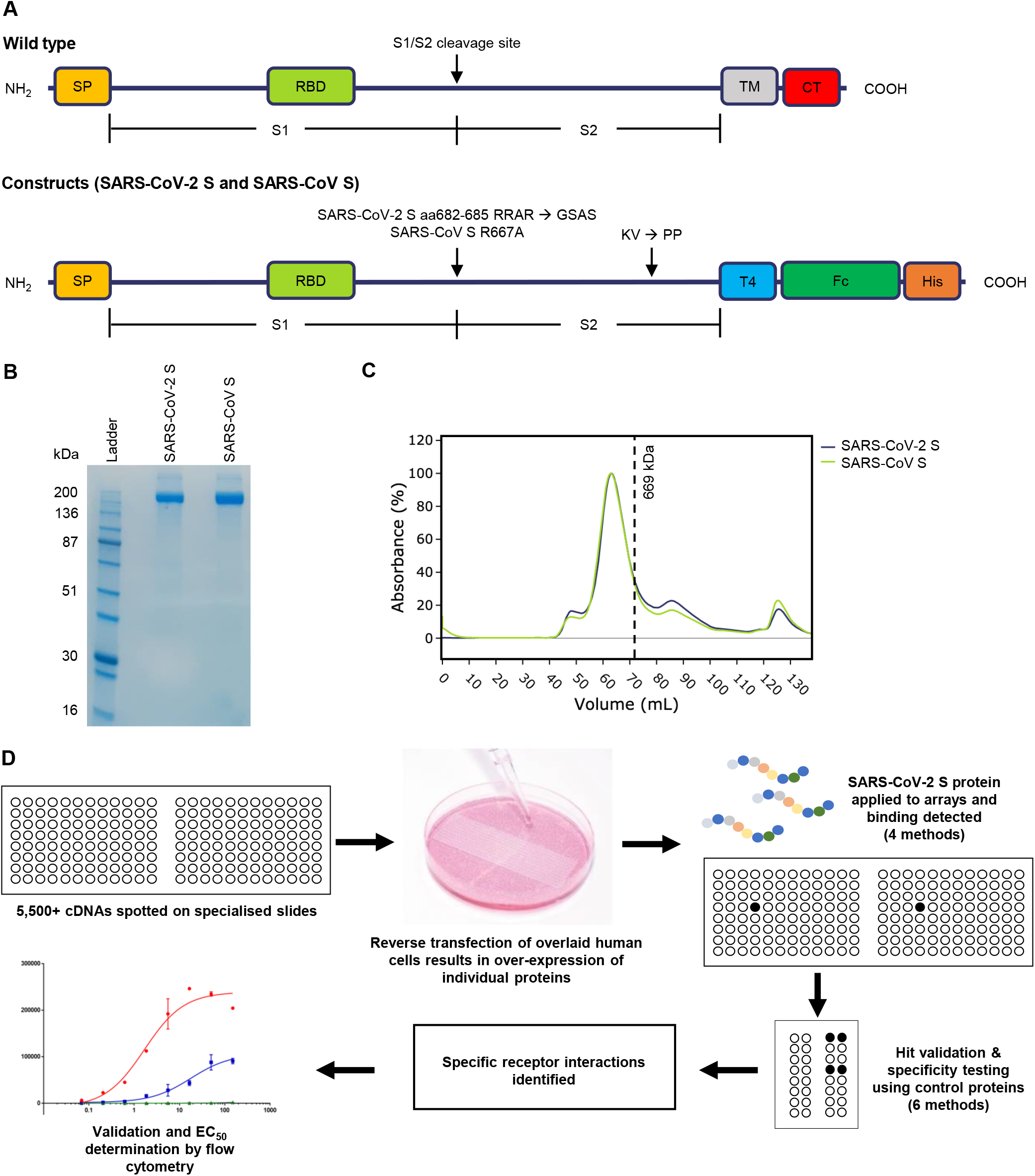
Overview of protein design and experimental receptor microarray approach. A. Schematic representation of spike protein constructs. SP, signal peptide; RBD, receptor binding domain; TM, transmembrane domain; CT, cytoplasmic domain; T4, T4 foldon trimerization domain; Fc, Fc region of human IgG; His, hexahistidine tag. B. Reducing SDS-PAGE analysis of purified recombinant spike proteins. C. Size exclusion chromatography UV absorbance trace of purified recombinant spike proteins, with retention volume of molecular weight standard thyroglobulin marked as a dashed line. D. Schematic of the experimental approach take to identify interactors of the SARS-CoV-2 spike protein using cell microarray-based expression library screens, followed by cell microarray and flow cytometric confirmatory screens.

The experimental approach to identify and validate interactors of the SARS-CoV-2 spike protein utilized systematic application of cell microarray-based expression library screens, followed by cell microarray and flow cytometric confirmatory screens, as outlined in Figure 1d.

### Identification of interactions for the SARS-CoV2 full-length spike protein

In order to identify receptors for SARS-CoV-2 spike protein, SARS-CoV-2 S protein was screened for binding to 5845 human HEK293 cell-expressed monomeric or heterodimeric human plasma membrane proteins and cell-surface tethered human secreted proteins using the Retrogenix cell microarray technology (Fig 1D). To maximize the likelihood of identifying interactions, full library screens were performed using 4 different approaches: SARS-CoV-2 S protein was added to live cell microarrays prior to cell fixation, or was added to fixed cell microarrays. In each case, slides were incubated with SARS-CoV-2 S protein followed by incubation with an AlexaFluor647 labelled anti-Fc detection antibody; or, SARS-CoV-2 S protein was pre-incubated with the same detection antibody at a 2:1 molar ratio prior to slide incubations.

Twenty-three positive hits were identified in total from the 4 library screens, with a full range of signal intensities. These comprised of 15 membrane proteins and 8 cell-surface tethered secreted proteins. All 23 hits, and 5 control proteins (CD20, CD86, EGFR and a FCGR3A:FCER1G heterodimer) were re-expressed in HEK293 cells on cell microarray slides, and a series of confirmatory screens were performed on fixed cells, live cells (before fixation) and live cells (no fixation) using the same SARS-CoV-2 S test protein, or SARS-CoV S control protein, each pre-incubated or sequentially incubated with the AlexaFluor647 detection antibody. Fig 2 shows a representative cell microarray confirmatory screen on live cells (no fixation) using the pre-incubation method. Interactions specific to SARS-CoV-2 S protein and SARS-CoV S control protein, that were not observed with the control treatments (Rituximab biosimilar and/or PBS) are shown in green (non-underlined); and interactions specific to SARS-CoV-2 S protein *but not* SARS-CoV S control protein, or the control treatments, are shown in green (bolded and underlined). The results from all cell microarray based confirmatory screens with the SARS-CoV-2 S test protein are summarised in Fig 3A, along with the library screen results with the same interactors. In total, 10 proteins were identified which were specific to SARS-CoV-2 S test protein, of which 5 were shared with SARS-CoV S protein (ACE2, ASGR1, CLEC4M, CD209, MSMP) and another 2 (CD44 and CLPS) were observed with MERS-S1 protein (Fig 3B). The 3 remaining interactors (CNTN1 and cell-surface tethered forms of the secreted proteins NID1 and APOA4) were exclusive to SARS-CoV-2 S test protein. Figs 3A and 3B show the relative intensity of each interaction, and the incubation methods that yielded positive hits. ACE2, CLEC4M and CD209 showed the highest signal intensities.

**Figure 2.**
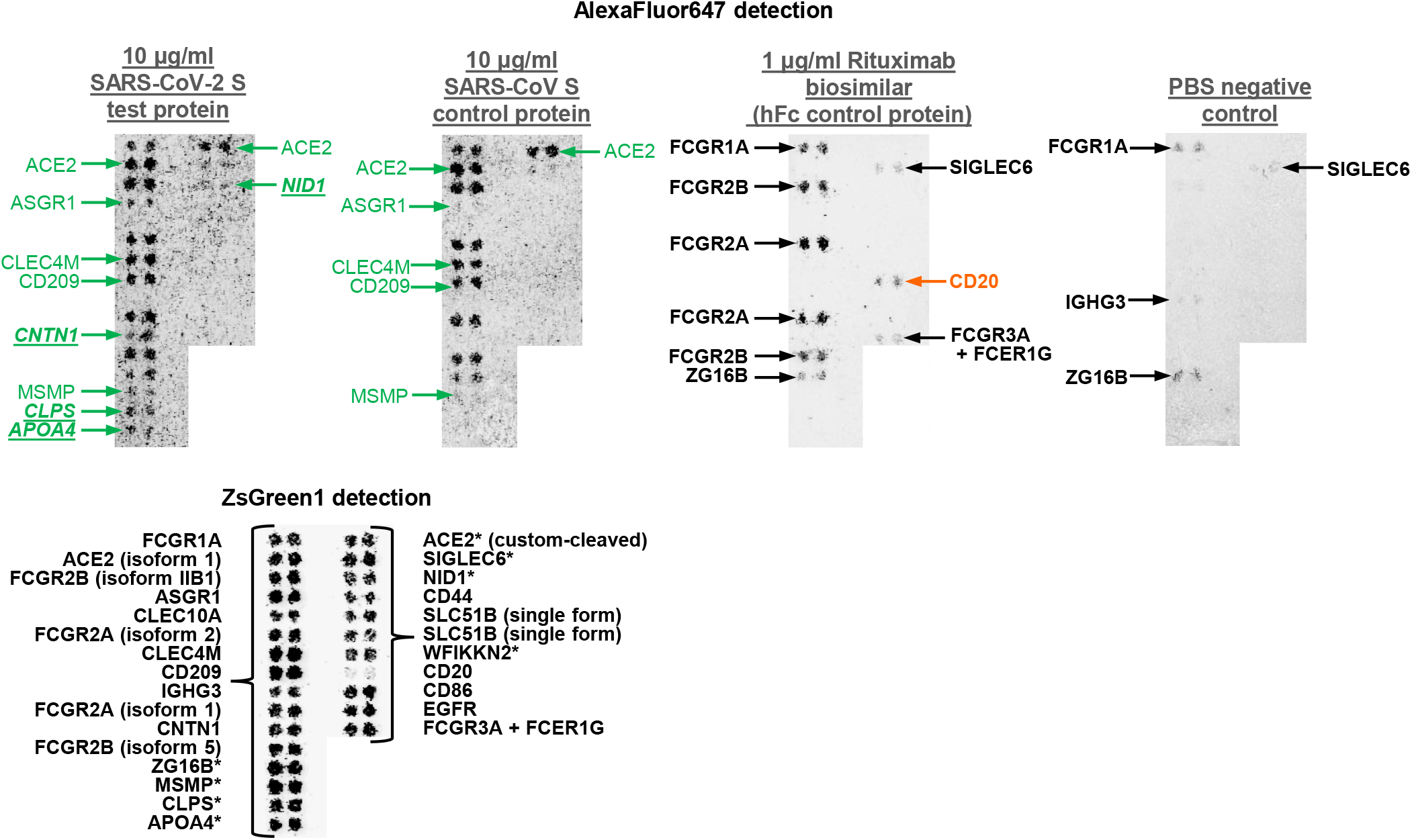
Confirmation, validation and comparison of cell microarray hits for SARS-CoV-2 and SARS-CoV. Representative images from one cell microarray-based confirmatory screen. Expression vectors encoding both ZsGreen 1 and individual library ‘hits’ were reverse-transfected into human HEK293 cells, and live transfectants were probed with the SARS-CoV-2 S test protein, SARS CoV S control protein, or other control treatments. This identified interactions that were specific to SARS-CoV-2 S test protein (bold and underlined), and those that were specific to SARS-CoV-2 S and SARS-CoV S spike proteins (green, not underlined). ZsGreen1 was used for spot localization. * indicates that these are secreted proteins which were synthetically cell surface-tethered.

**Figure 3.**
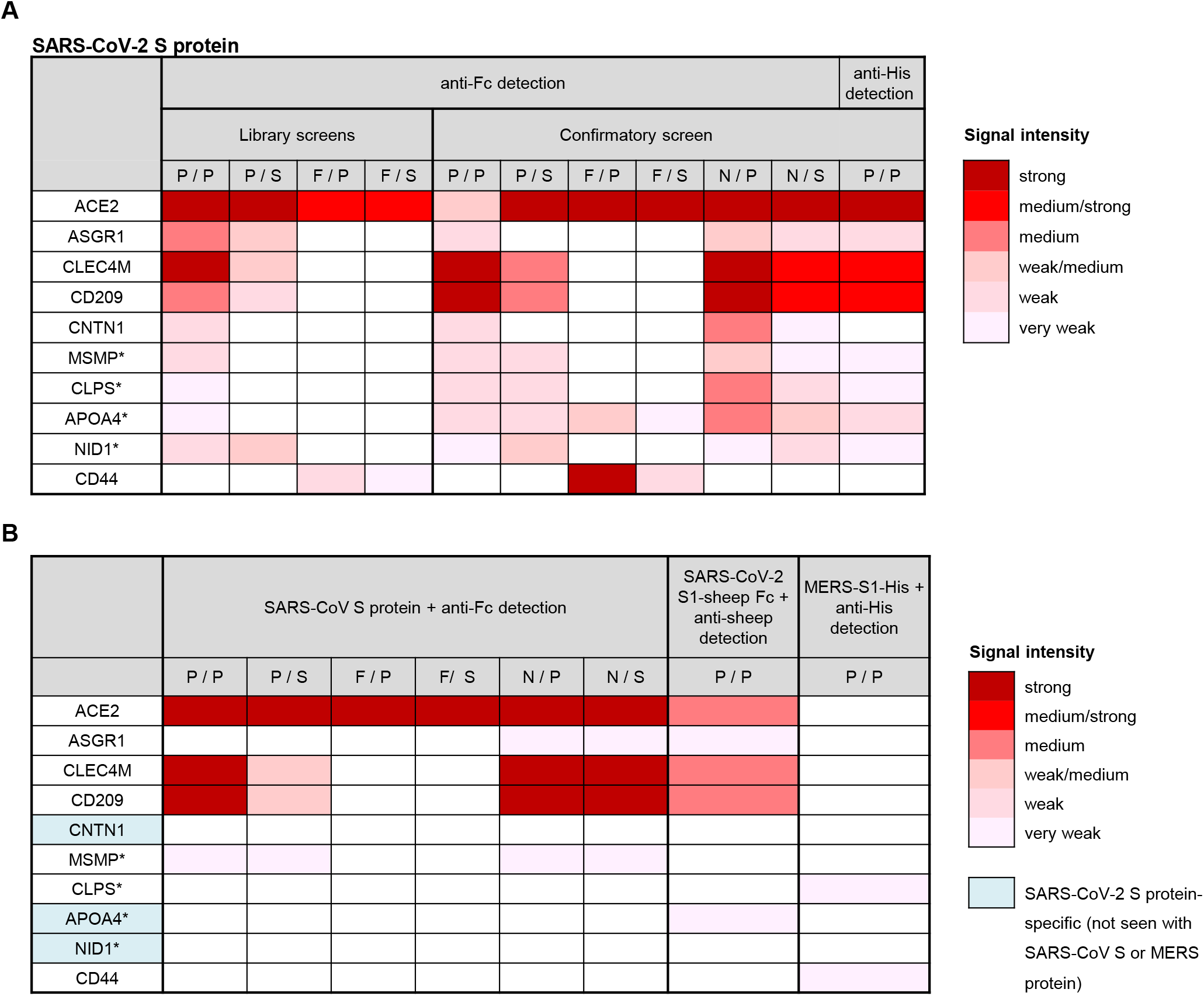
Specificity of SARS-CoV-2 binding interactions. A. Representative of all cell microarray screening data arising library screens with 10 μg/mL SARS-CoV-2 S test protein using 4 different incubation approaches, and from the cell microarray-based confirmatory screens using 6 different incubation approaches. Either a fluorescent anti-Fc detection antibody or a fluorescent anti-His detection antibody was used, as indicated. Interaction intensities against the 10 SARS-CoV-2 S test protein-specific hits were graded from very weak to strong intensity. White boxes indicate no interaction was observed. * indicates that these are secreted proteins which were cell surface-tethered. B. The 10 SARS-CoV-2 S test protein-specific hits were tested for cross-reactivity with three control proteins, 10 μg/mL SARS-CoV S protein (using anti-Fc detection), 5 μg/mL SARS-COV-2 S1 fragment-sheep Fc fusion protein (using an anti-sheep detection antibody) or 5 μg/mL His-tagged MERS-S1 protein (using an anti-His detection antibody). Interaction intensities were graded from very weak to strong intensity. White boxes indicate no interaction was observed. P/ means added prior to fixation, F/ means added after fixation, N/ means no fixation, /P means a control protein was pre-incubated with the detection antibody at a 2:1 molar ratio, and /S means a control protein was incubated with the cells first, followed by the sequential incubation with detection antibody. Interactions exclusive to SARS-CoV-2 are highlighted in blue.

Five of the 10 interactions (ACE2, ASGR1, CLEC4M, CD209 and APOA4) were also detected using a commercial SARS S1- sheep Fc domain fusion protein (Fig 3B). Nine of the 10 interactions (the exception being CNTN1) were detected when binding of the SARS-CoV-2 S test protein binding was detected using an AlexaFluor647 anti-His detection instead of AlexaFluor647 anti-hIgG Fc detection (Fig 3A). With both detection approaches, ACE2, CLEC4M and CD209 showed the highest signal intensities. The fact that a control MERS-S1-His protein showed detectable binding to CLPS and CD44 only indicates that the other interactions are not general to all coronaviruses.

### Validation of the identified SARS-CoV-2 spike protein interactions

In order to investigate the SARS-CoV-2 spike protein interactions further, nine of the 10 interactions identified above (ACE2, ASGR1, CLEC4M, CD209, CNTN1, MSMP, APOA4, NID1 and CD44) were investigated further on live transiently transfected HEK293 cells by flow cytometry, using a concentration range of 0–150 μg/mL of SARS-CoV-2 S test protein or SARS-CoV S control protein. The exception was CLPS, since an initial single dose experiment showed this interaction could not be reproduced by flow cytometry. Fig 4 shows titration curves of the two spike proteins against each expressed target, after normalising for background binding to cells at each spike protein concentration. EC50 values were determined where possible, and are independent of the relative expression level of each target protein (Table 1). These data confirmed the specific binding of SARS-CoV-2 S protein to each of the 9 identified target proteins, with EC50 values for binding ranging from 1.6 ± 0.5 μg/mL (ACE2) to greater than 150 μg/mL (CNTN1 and CD44, for which EC50 values could not be determined using the concentrations tested). For 5 of the target proteins, namely ACE2, ASGR1, CLEC4M, CD209 and MSMP, there was no significant difference in the EC50 values between the SARS-CoV-2 and SARS-CoV S proteins. In contrast, for 3 of the target proteins (CNTN1, APOA4 and NID1), while no significant binding of SARS-CoV S protein was detected, SARS-CoV-2 S protein showed specific binding interactions, consistent with the cell microarray data (Figs 3A and 3B). Notably, SARS-CoV-2 S protein interacted with APOA4 as tightly as its interaction with ACE2 (EC50 values: 1.8 ± 0.5 μg/mL and 1.6 ± 0.5 μg/mL, respectively).

**Figure 4.**
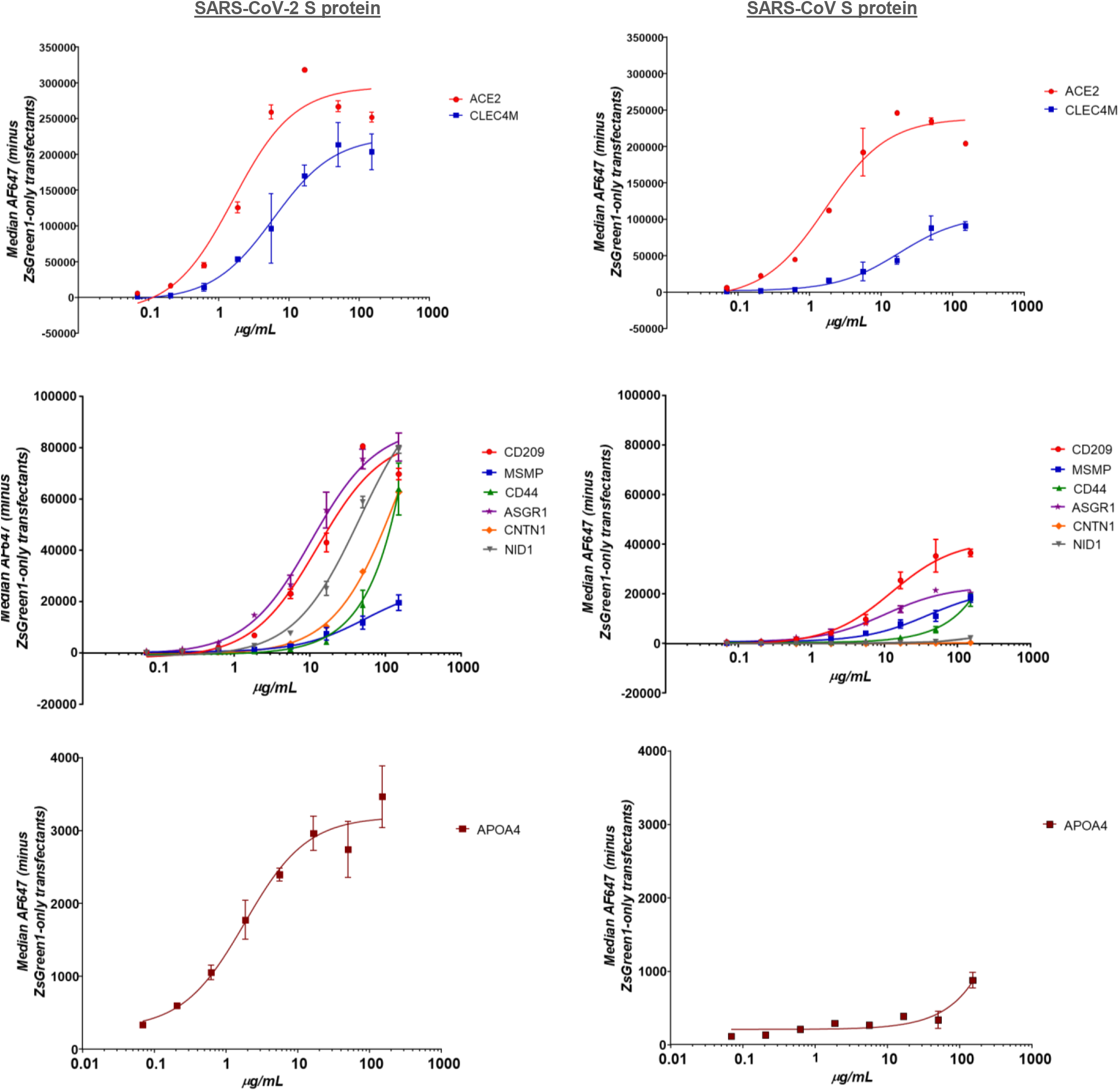
Flow Cytometric analysis of binding of cell surface receptors between SARS-CoV-2 and SARS-CoV. Live human HEK293 cells transiently over-expressing both ZsGreen and each specific hit identified by cell microarray, were incubated with 0 to 150 μg/mL of SARS-CoV-2 S protein (left hand panels) or SARS-CoV S protein (right hand panels), and interactions were investigated by flow cytometry. After correcting for background binding, binding curves were generated against each target, from which EC50 values were calculated.

**Table 1.** Unique binding interactions of SARS-CoV-2. Table of EC50 values (± standard errors) for binding of SARS-CoV-2 S protein (left) or SARS-CoV S protein (right), against each of the 10 specific protein interactors. Binding interactions were similar between SARS-CoV-2 S protein or SARS-CoV S protein, except for CNTN1, APOA4 and NID1, which were exclusive to SARS-CoV-2 S protein. The EC50 value for binding of SARS-CoV-2 S protein to APOA4 was comparable to that with ACE2.

|  | SARS-CoV-2 S protein | SARS-CoV S protein |
| --- | --- | --- |
|  | EC<sub>50</sub> (µg/mL) | EC<sub>50</sub> (µg/mL) |
| ACE2 | 1.6 ± 0.5 | 1.7 ± 0.5 |
| ASGR1 | 10.5 ± 1.4 | 10.2 ± 1.9 |
| CLEC4M | 6 ± 1.6 | 17.9 ± 5.1 |
| CD209 | 12.7 ± 3.3 | 12.5 ± 2.8 |
| CNTN1 | >150 | ND |
| MSMP | 54.5 ± 15.9 | 39.6 ± 10.5 |
| CLPS | NI | NI |
| APOA4 | 1.8 ± 0.5 | ND |
| NID1 | 45.4 ± 5.3 | ND |
| CD44 | ND | ND |

ND = Not Determinable at dose range tested (0 - 150 µg/mL)
NI = Not Investigated

## DISCUSSION

In this study we employed an agnostic approach to the identification of human plasma membrane and secreted protein binding partners for the SARS-CoV-2 S protein, in order to identify potential therapeutic targets for viral cell entry. We screened full length SARS-CoV-2 spike trimer protein against a cell microarray platform, expressing a vast library of membrane and secreted proteins expressed in human cells. We employed four different incubation methodologies to maximise the success rate. The value of this platform technology is demonstrated by the confirmation of a number of previously known receptors as well as the discovery of three novel receptors.

The sequence of the SARS-CoV-2 was available within 1 month of the initial cases of the COVID pandemic being reported in Wuhan in Dec 2019 (Wu et al, 2020). This showed the virus shared structural similarities with SARS-CoV exhibiting a spike protein on the surface of the virion responsible for receptor binding and fusion, that is heavily glycosylated and formed by 3 S protein monomers. In our study, we engineered the SARS-CoV-2 S sequence to produce full-length S monomers that assembled to form a homo-trimer, ensuring the S protein was in its physiologically relevant prefusion conformation. This enabled us to look solely at initial binding events.

The unique cell microarray platform uses an array of expression vectors encoding a total of more than 5,800 full length human integral plasma membrane and membrane-tethered secreted proteins, which are reverse transfected and individually over-expressed on the cell surface of human HEK293 cells. This physiologically-relevant system allows even low affinity interactions with the viral spike protein to be detected with a high degree of sensitivity and specificity.

Using this technology platform, we confirmed the binding of SARS-CoV and SARS-CoV-2 spike proteins to ACE2 with equivalent affinities (half-maximal binding was achieved at 1.7μg/ml (2.8nM) and 1.6μg/ml (2.7nM) respectively). We also confirmed binding of SARS-CoV-2 S at other receptors previously reported, namely ASGR1 (CLEC4H1), CLEC4M and CD209 (CLEC4L) (Gu et al, 2020; Brufsky et al, 2020).

Importantly three novel binding proteins, APOA4, CNTN1 and NID1 were identified which showed specificity for SARS-CoV-2 when compared with SARS-CoV. Notably, SARS-CoV-2 S protein demonstrated an equivalent affinity for APOA4 as for ACE2 (half-maximal binding was achieved at 1.8μg/ml (3nM) and 1.6μg/ml (2.7nM) respectively).

While we had speculated alternative receptors or co-receptors of SARS-CoV-2 might exist in addition to ACE2 this is the first time APOA4 has been identified, suggesting this is another mechanism by which SARS-CoV-2 may infect cells. APOA4 is a lipoprotein, previously shown to influence viral attachment and interaction of Hepatitis C virus with its primary receptors (Zeisel et al, 2013). APOA4 is expressed in the intestine, is a cellular marker of enterocytes and plays role in lipid biosynthesis and metabolism. Expression of APOA4 also correlates with the highest level of ACE2 expression, which is found in intestinal enterocytes (Hamming et al, 2004; Qi et al, 2020).The gastrointestinal tract is considered a target organ for SARS-CoV-2 infection with up to 34% of COVID-19 patients reporting digestive symptoms like diarrhea, nausea, and abdominal discomfort (Yang & Tu, 2020). While this may suggest a link for APOA4 and SARS-CoV-2 infection in the gastrointestinal tract, further work is required to determine its functional significance.

Our study also confirmed a number of receptors belonging to the C-Type Lectin Domain Family 4 (CLEC4), which exhibit high avidity for glycoproteins, that are present in viral envelope proteins (Helenius et al, 2018; Feinberg et al, 2001) and also the SARS-CoV-2 S protein. Of significance, CD209 (CLEC4L/ DC-SIGN) has been cited as a coreceptor for SARS-CoV where deglycosylation of the spike protein reduces infectivity of viral pseudotypes expressing SARS-CoV S. Mutations in the SARS-CoV-2 S protein, which increase glycosylation were identified in clinical isolates, and shown to increase viral load and cytopathic effects in culture (Han et al, 2007; Yao et al, 2020). Interestingly, as viral glycosylation occurs in the host cell endoplasmic reticulum and Golgi apparatus, it is subject to host specific and genetic influences, and hence the family of CLEC4 proteins offer potential for effect on virulence in different ethnicities.

Lectins differ in tissue expression and ligand affinities and therefore could influence the tissue specific tropism seen in SARS-CoV-2 infection and COVID-19 disease pathology. Especially of interest are those lectins showing tissue specific expression in organs associated with disease pathology and where ACE2 expression is low. ASGR1 (CLEC4H1) is an endocytic recycling receptor which has been reported to facilitate entry of Hepatitis C virus (Saunier et al, 2003) and is expressed exclusively in the liver. ACE2 however is expressed at low levels in the liver, yet liver impairment is common in patients with COVID-19, where SARS-CoV-2 infection directly causes cytopathy of hepatocytes and impairs liver function.

Overall, our data suggests COVID-19 may be a highly promiscuous virus exploiting multiple mechanisms of cell entry, contributing to its severity and targeting of specific patient groups. Furthermore, using recombinant full length spike trimer protein coupled with cell microarray technology and flow cytometry, we have demonstrated a fast, accurate and comprehensive approach for discovering the human cell surface targets of viruses. This technology can be further applied to this pandemic, with the many emerging variants of SARS-CoV-2, and to future epidemics for the identification of exploitable drug targets to control viral infection.

## MATERIALS & METHODS

### Construct design, protein expression and purification

Constructs encoding SARS-CoV-2 spike protein aa14-1213 and SARS-CoV spike protein aa14-1195 were synthesised commercially codon optimised for mammalian cell expression (Genscript and GeneArt, respectively). Residues 682-685 were mutated from RRAR to GSAS in SARS-CoV-2 S and R667A mutation was introduced in SARS-CoV S, as well as two stabilising mutations (K986P and V987P in SARS-CoV-2 S, K968P and V969P in SARS-CoV S) and a T4 foldon trimerization domain at the C-terminus. Both spike proteins were also fused to the Fc region of human IgG for assay detection and to a hexahistidine tag for purification at the C-terminus. The resultant recombinant proteins, SARS-CoV-2 S and SARS-CoV S respectively, were produced in the HEK-293 cell system and were purified from the culture supernatants by capture with Ni Sepharose excel affinity resin (Cytiva) followed by size exclusion chromatography using a Superose 6 16/60 column in phosphate-buffered saline (PBS). The purified proteins were analysed by reducing SDS-PAGE for purity, mass spectrometry for identity and A280 correcting for the extinction coefficient in order to determine concentration.

### Cell microarray library screening

5845 mammalian cell expression vectors, each encoding a full-length human plasma membrane protein or a cell-surface tethered secreted protein, including ACE2, were arrayed in duplicates across 16 slide sets. Human HEK293 cells were grown over the vector microarray, leading to reverse transfection at each location. ZsGreen1 encoded by the vector was used to monitor the transfection efficiency and define the microarray position. All transfection efficiencies exceeded the minimum threshold.

Either before or after fixing the cells, 10 μg/mL SARS-CoV-2 S protein alone, or pre-incubated with AlexaFluor647-conjugated goat anti-human IgG Fcγ pAb (Jackson ImmunoResearch; detection antibody) at a 2:1 molar ratio (‘pre-incubation method’), was incubated with the cell microarray slides for 1 hour. In the case of cell microarray slides incubated with SARS-CoV-2 S protein alone, the same detection antibody was added sequentially (‘sequential incubation method’). Library hits (duplicate Alexa-Fluor647-positive spots) were captured by fluorescence imaging (GE Ettan DIGE imager), and analysed using ImageQuant software (GE Healthcare).

### Cell microarray confirmatory and specificity screening

Vectors encoding all library hits and negative control proteins (CD20, CD86 and EGFR) were arrayed in duplicate, and human HEK293 cells were reverse-transfected as described above. Cell microarray slides were incubated before, after, or in the absence of cell fixation with 10 μg/mL SARS-CoV-2 S test protein, 10 μg/mL SARS-CoV S control protein, 1 μg/mL Rituximab biosimilar (human IgG1; Absolute Antibody, UK), or phosphate-buffered saline (PBS) only, each pre-incubated with AlexaFluor647-conjugated goat anti-human IgG Fcγ pAb at a 2:1 molar ratio, or treated sequentially with the same detection antibody. Identical slides were treated before cell fixation with 5 μg/mL SARS-CoV-2 S1-sheep Fc (Native Antigen Company, UK, catalogue # REC31806) or PBS, each pre-incubated at a 2:1 molar ratio with AlexaFluor647 donkey anti-sheep IgG (H+L) (Thermo Fisher, UK, catalogue # A-21448); or 5 μg/mL SARS-CoV-2 S test protein (Peak Proteins, UK, described above), 5 μg/mL MERS S1-His (Native Antigen Company, UK, catalogue # REC31760) or PBS, each pre-incubated at a 2:1 molar ratio with Penta His Alexa Fluor 647 Conjugate (Qiagen, UK). Hits (duplicate AF647 positive spots) were captured and analysed as described above.

### Hit validation by flow cytometry

Expression vectors encoding ZsGreen1 alone, or encoding ZsGreen1 and one of 9 cell microarray hits were transfected into human HEK293 cells. After 48 h, cells were treated with Accutase (Sigma-Aldrich, UK), pelleted by centrifugation, and resuspended into incubation buffer (PBS supplemented with 10% foetal calf serum) containing a dose range of 0 to 150 μg/mL SARS-CoV-2 S protein or SARS-CoV S protein, or 1 μg/mL Rituximab biosimilar (human IgG1; Absolute Antibody, UK). After a 1 hour incubation, cells were again pelleted, washed with PBS, and resuspended into incubation buffer containing AlexaFluor647-conjugated goat anti-human IgG Fcγ pAb for a further incubation. Cells were further washed with PBS, and analysed by flow cytometry using an Accuri (Becton Dickinson, UK). A 7-Aminoactinomycin D live/dead dye was used to exclude dead cells in the analysis, and ZsGreen-positive (transfected) cells were selected for analysis.

For each target transfectant, at each concentration of SARS-CoV-2 S protein or SARS-CoV S protein, the median AlexaFluor647 value was corrected for background binding to the cells by subtracting median binding to ZsGreen-only transfected cells at the same concentration. Dose titration curves were plotted, and EC50 values determined, using Prism GraphPad.

## AUTHOR CONTRIBUTIONS

Conceptualisation and study design LRR, SB, JF, JS, MA; Protein production RFR, CG, MA; Investigation HT, JD, LC; Data analysis RFR, MA, JF, JS; Writing the manuscript SB, JF, RFR LRR; Provided conceptual guidance, support and review LRR, SB, MA

